# Biogenic methane cycling is controlled by microbial cohorts

**DOI:** 10.1101/2022.01.24.477581

**Authors:** Rhiannon Mondav, Gaëtan Martin, Sari Peura, Sarahi L. Garcia

## Abstract

The generation and consumption of methane by aquatic microbial communities is an important contribution to the global carbon budget. We sought to broaden understanding of consortia members and interactions by combining multiple methods including analysis of natural and cultivated microbial communities. By analysing the microbial community composition and co-occurrence patterns of a lake time-series we were able to identify potential consortia involved in methane cycling. In combination with methane flux, we also analysed the community composition and co-occurrence patterns of reduced microbial model communities with inoculum from the same lake. While the network analyses confirmed many known associations, when combined with results from mixed cultures, we noted new players in methane cycling. Cultivated model communities were shown to be an effective method to explore the rarer but still important players in methane cycling and for identifying new putative members. Here we show that using multiple methods to approach the complex problem of methane cycling consortia yields not just insights into the known taxa but highlights potential new members creating new hypotheses to be tested.

## Introduction

Methane emissions from biogenic sources are determined by the ratio and activity of methanogenic microbes producing and methanotrophic microbes consuming this powerful greenhouse gas. Considering that methane is responsible for 17% to 20% of the radiative forcing component of global warming (1, 2), elucidating all players in the methane cycle involved in the net methane emissions from various biogenic sources is of utmost importance. It is known that methanotrophs can oxidize between 30 and 99% of CH4 produced in the sediments and the water column before it reaches the atmosphere (3–6). However, the detection of methanotrophs in an environment does not necessarily correlate with methanotrophic activity, and other factors may determine whether methane is oxidized (7).

One factor that may affect the rate or existence of methanotrophy is the composition of the microbial community. For example, it has been shown that aerobic methanotrophs can be active in anoxic waters when co-occurring with photosynthetic organisms (8), suggesting that the methanotrophs utilize oxygen liberated by these phototrophs for methane oxidation. However, discovering and pinpointing more of these symbiotic interactions is difficult in natural environments as the number of possible interacting organisms is very high. Methods to cultivate communities with reduced diversity have been developed (9) to overcome this limitation and study organisms in a model communities in conditions mimicking their natural habitat. In this method, environmental samples are diluted in filtered water from their own environment and microorganisms subsampled for cultivation. These so-called model communities are a powerful way to study the microbial interaction in a semi-natural environment (10). Model communities also provide tools for building hypotheses that could then be further tested by looking into interaction networks in time series of natural environments (11).

For this study, we established 177 dilution-cultures model communities using samples from Lake Lomtjärnan, a lake in Sweden with high methane concentrations in the water column to examine the methane cycling capacity of microbial assemblages. We also utilized a 2-week depth discrete timeseries taken from March to April 2016 to examine the network around the methane oxidizing communities in the actual lake. We hypothesize that the capacity to produce or oxidize methane is not only related to the presence of organisms with the known ability to produce or use methane, but that their cohorts might be key controllers in methane cycling.

## Materials and methods

### Lake water collection and media preparation

Environmental samples for the time-depth-series were collected and published previously (12). In brief, a sampling campaign including six time points was done in winter the last week of March and the first week of April 2016 on Lake Lomtjärnan. This small forest lake had an ice cover at the time of sampling and is located in central west region (Jämtland) of Sweden (Fig 1A). The surface area of the lake is about 1 ha, and the maximum depth is 3.5 m. The lake is located on a mire surrounded by a coniferous forest. At each sampling occasion, samples at six different depths (0.65 m, 1.0 m, 1.35 m, 1.85 m, 2.35 m, 2.75 m) were taken to create a time-depth-series of the lake totalling 35 samples. The deepest depth was not taken on the first sampling because samples were taken in a location where the max depth was around 2.35 m.

**Figure 1.**
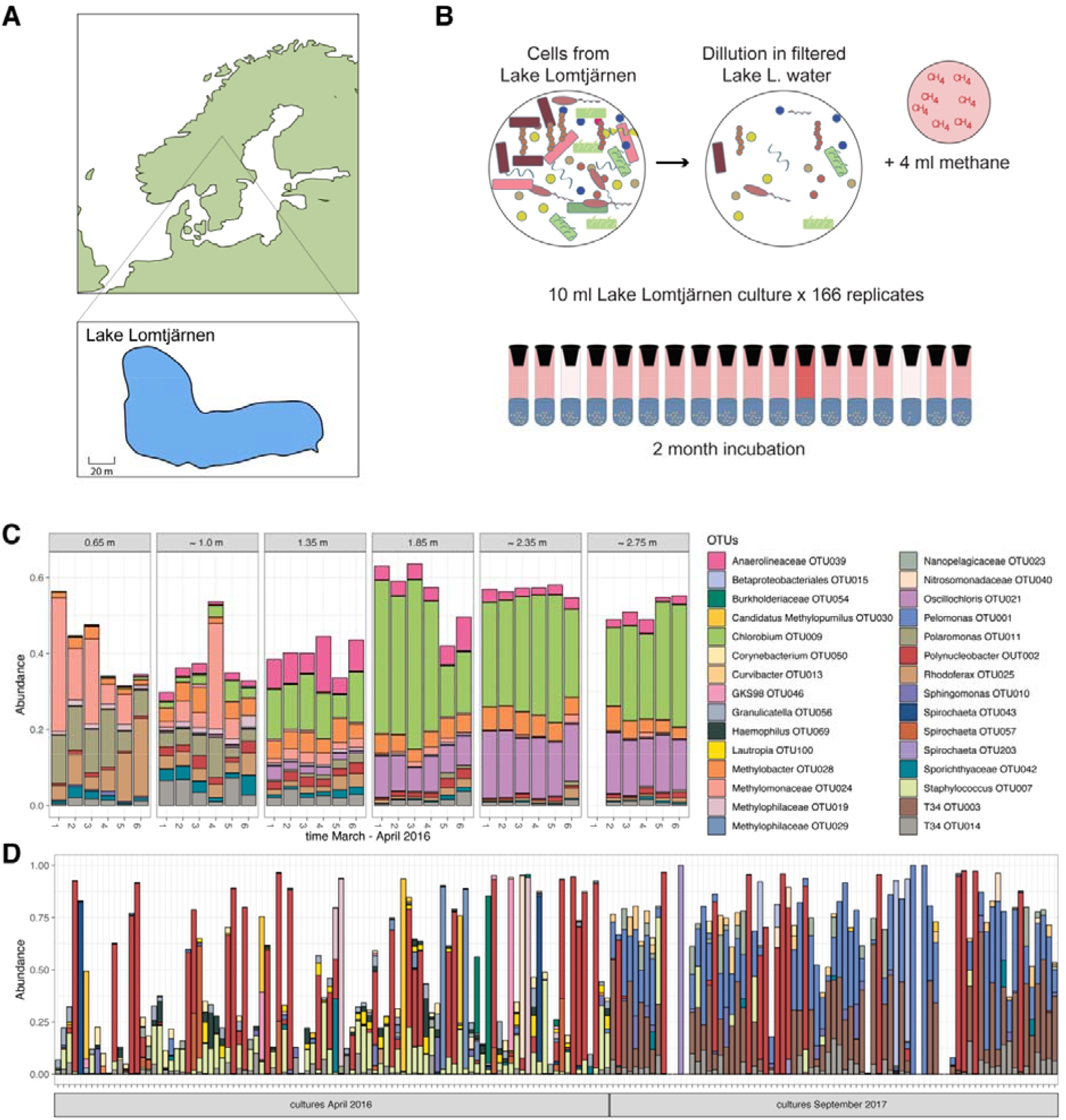
Sample location [A], schematic of dilution and set up of model communities [B], most abundant OTUs detected in lake timeseries [C], and most abundant OTUs from model communities [D]. The legend in C is also for D.

Water for the time-depth-series was collected using a depth-discrete Limnos tube-sampler (Limnos, Poland) and the water was subsequently filtered through a Sterivex filter (0.22 μm) and the filters were stored immediately in liquid nitrogen for later DNA extraction. Water to be used for both growth media and inoculum was collected from the anoxic layer on the sixth sampling occasion of the lake in April 2016 and again in September 2017.

The water for media was collected using the same depth-discrete Limnos tube-sampler (Limnos, Poland) and was quickly poured into 1 litre Schott bottles which were filled and let to overflown to remove all the oxygenic water prior to closing the bottles. These bottles were then kept in the dark at 4°C for 2 days to ensure anoxia. The water for media was filtered twice through 0.2□μm Sterivex filters (Millipore) inside the anaerobic glove box, and the filtered-sterilized media was collected in a sterile Schott bottle and closed. These bottles were further exposed to UV light for 10 min.

The inoculum for the cultures was collected using the Limnos tube-sampler in a falcon tube and was then flash frozen with liquid nitrogen and kept in -80 °C until the cultures were established. The following steps of the preparation were performed under anoxic atmosphere using an anaerobic glove box.

### Preparation of serum bottles, cell inoculant and cultures

20 ml serum bottles, crimps, and stoppers were autoclaved at 120°C for 20 min. To limit the risk of potential bactericides such as benzyltoluenes and phenylalkanes, the stoppers were autoclaved submerged in deionized water, and subsequently rinsed and boiled in sterile deionized water and finally cooled in sterile deionized water. After autoclaving, the serum bottles were sealed with these sterilised and detoxified butyl rubber stoppers and aluminium crimps. The incubation vessels were assembled under a laminar flow, using autoclaved tools, before being flushed at least 3 times with nitrogen to remove O2. To maintain the sterility of the bottles during the flushing operation, new sterile needles were used for each bottle and the stoppers were carefully disinfected with 90% ethanol. Furthermore a 0.2 μm filter was used to prevent contamination from the N2 gas flow.

The sealed sterile and anaerobic serum bottles were supplemented with 10 ml of media, a.k.a. filtered lake water. Subsamples of the unfiltered lake water from Lomtjärnan April 2016 (average 5×106 cells/ml) and September 2017 were run through a flow cytometer to estimate cell concentration. Based on those values, another subsample, not exposed to the atmosphere was diluted with lake-water-media to approximatively 50 cells/ml. To inoculate the bottles, 1 ml of this 50 cells/ml solution was then injected using sterile needles and syringes into the 20 ml serum bottles. The number of cultures prepared were: 98 from 2016 and 79 from 2017. A few bottles were incubated without inoculum as media control cultures. To test for the potential for anoxic methane oxidation 4 ml of a mix of CH4 and CO2 with a ratio of 80/20 % were injected into each bottle (Figure 1B).

Bottles were then incubated at 11°C for two months in the dark, to mimic lake conditions, for the samples taken in the autumn and under dim light for the samples collected in the spring. After the two-month incubation, 1 ml of the headspaces was sampled for gas analysis. Methane content was measured using the gas analyser Biogas 5000 (Geotechnical Instruments, UK) and gas chromatography (Clarus 500, Perkin Elmer, USA, Polyimide Uncoated capillary column 5m x 0.32mm, FID detector). At the same time as gas sampling 200 μl of the cultures were collected and preserved at -80°C for DNA analysis.

### 16S rRNA gene amplicon preparation and sequencing

DNA was amplified directly from 1 μl of culture. Library preparation for 16S rRNA gene analysis was done following a two-steps Polymerase Chain Reaction (PCR) protocol, as described in Mondav et al 2020 (11). All PCRs were conducted in 20 μl of volume using 0.02 U/μl Phusion high fidelity DNA polymerase, 1X Q5 reaction buffer (NEB, UK), 0.25 μM primers and 200 μM dNTP mix and 1 μl mixed culture template. The first step was performed in triplicate with primers 341F (3’-CCTACGGGNGGCWGCAG-5’) and 805NR (3’-GACTACNVGGGTATCTAA-5’) (13). The thermal program consisted of 20 cycles with an initial 98°C denaturation step for 10 min, a cycling program of 98°C for 10 seconds, 48°C for 30 seconds, and 72°C for 30 seconds and a final elongation step at 72°C for 2 minutes. Triplicate PCR reactions were then pooled and purified with magnetic beads (Sera-Mag™ Select, GE Healthcare, Chicago, United States of America), and 2 μl of the purified products were used at a template for a second stage PCR, where indexed primers were added. The second thermal program consisted of 15 cycles with an initial 98°C denaturation step for 30 seconds, a cycling program of 98°C for 10 seconds, 66°C for 30 seconds, and 72°C for 30 seconds and a final elongation step at 72°C for 2 minutes. Following amplification, PCR products were again purified with magnetic beads and quantified with Qubit™ using the Qubit™ dsDNA HS Assay Kit (Invitrogen™). Finally, 15.6 μg of each indexed and purified PCR product were pooled before submission of the sample to the Science for Life Laboratory SNP/SEQ sequencing facility hosted by Uppsala University (Uppsala, Sweden). Sequencing was done using Illumina Miseq in paired-end mode with 300bp and v3 chemistry.

### Amplicon bioinformatics

Sequence processing was performed with Mothur 1.41.0 following the MiSeq SOP (14), with the exception that clustering to operational taxonomic units (OTUs) was done using VSEARCH (15) as implemented in Mothur (16). Taxonomy was assigned against the Silva 132 database (17). Amplicon sets were separated into groups of sample or inoculant origin: time-depth-series lake communities, anaerobic lake cultures 2016, anaerobic lake cultures 2017, and 2 un-inoculated media controls. Nine lake-2017 cultures produced only a few sequences (<20) and were removed from the dataset. Remaining sequences were then subsampled once using Qiimes (18) single rarefaction to an even depth of 600 reads per sample for the un-inoculated media, and 2000 reads per sample for all others. This resulted in final sample counts: 35 lake time-depth-series, 93 lake-2016 cultures and 70 lake-2017 cultures. These normalized OTU tables were used for compositional and comparative analyses. Column graphs of the ten most abundant phylotypes from each dataset were made for compositional analyses. OTU tables were processed and visualised in R 4.0.3 (19) using phyloseq 1.34.0 (20) and ggplot2 (21).

Prior to co-occurrence analysis for network visualisation, OTUs present in less than 10% of each culture group, 50% of the lake water samples, or with a relative abundance always below 1 % were removed. This was done to reduce sparcity to below 50%. A network ensemble approach was used where any correlation (edge) between OTUs (node) that was found in at least two of the three following methods was included in the final visualisation: SPIEC-EASI (22), SparcCC (23), and Pearsons correlation. All co-occurrence methods were implemented in R 4.0.3 using spiec-easi 1.1.1 for SPIEC-EASI, SparcCC and the psych 2.1.3 package for Pearsons correlation. Networks were visualised in cytoscape.

To identify communities that had unbalanced methane cycling (flux), all cultures that had a recorded change in methane concentration and all cultures that had at least one phylotype associated with methane generation or consumption were retained and analysed for OTU co-occurrence.

### Data availability and source code

All scripts used to calculate network correlations are available at: https://github.com/rmondav/methane_cycling_networks, v1.0.0 recorded at https://doi.org/10.5281/zenodo.5531947. Time-depth-series amplicons are available under BioProject PRJEB27633 (12). Mixed culture sequence data has been deposited at the European Nucleotide Archive (ENA) at EMBL-EBI under accession number BioProject PRJEB48661 (https://www.ebi.ac.uk/ena/browser/view/PRJEB48661).

## Results

### Change in dominant lake community with depth

The most abundant OTUs in the lake time-depth-series belonged to just three phyla: Bacteroidetes, Chloroflexi, and Proteobacteria. The surface lake microbial community was dominated by the □-proteobacterial Rhodoferax OTU25 and Polaromonas OTU11 and Methylomonaceae OTU24 while the deeper lake community was increasingly dominated by a Chlorobium OTU9 and Oscillochlorus OTU21 (Fig. 1C). Polynucleobacter OTU2 was highly abundant throughout the lake water (Fig 1C) showing no preference for depth. It was also abundant in most of the mixed cultures (Fig 1D) and was present at low abundance in media controls from Sept 2017 (0.2 μm filterable microbes).

### Communities in lakes associated with methane cycling

Known methanotrophs (obligate methane consumers) in the lake time-depth-series network were associated with either a phototrophic cluster of bacteria (green circle top left) or clustered around a methanogen (gold circle top centre, Figure 2). Carbon fixing phototrophs as detected in this cluster, methanotrophs, and the methanogen are all obligate C1-carbon molecule cyclers. Sulphate-reducing bacteria were also associated with these two clusters. The Oscillochloris OTU21 and Chlorobium OTU9 were dominant phylotypes of deeper lake (green circle Figure 2). The methanotroph Methylomonaceae OTU24 and Polaramonas OTU11 (grey circle bottom Figure 2) were dominant in lake Lomtjärnan surface waters (Fig 1B). This cluster (grey circle) was negatively correlated to the other methanotrophs, the phototrophs and the methanogen. Methylotrophs (facultative methane consumers) had no clear association with any of these metabolic clusters.

**Figure 2.**
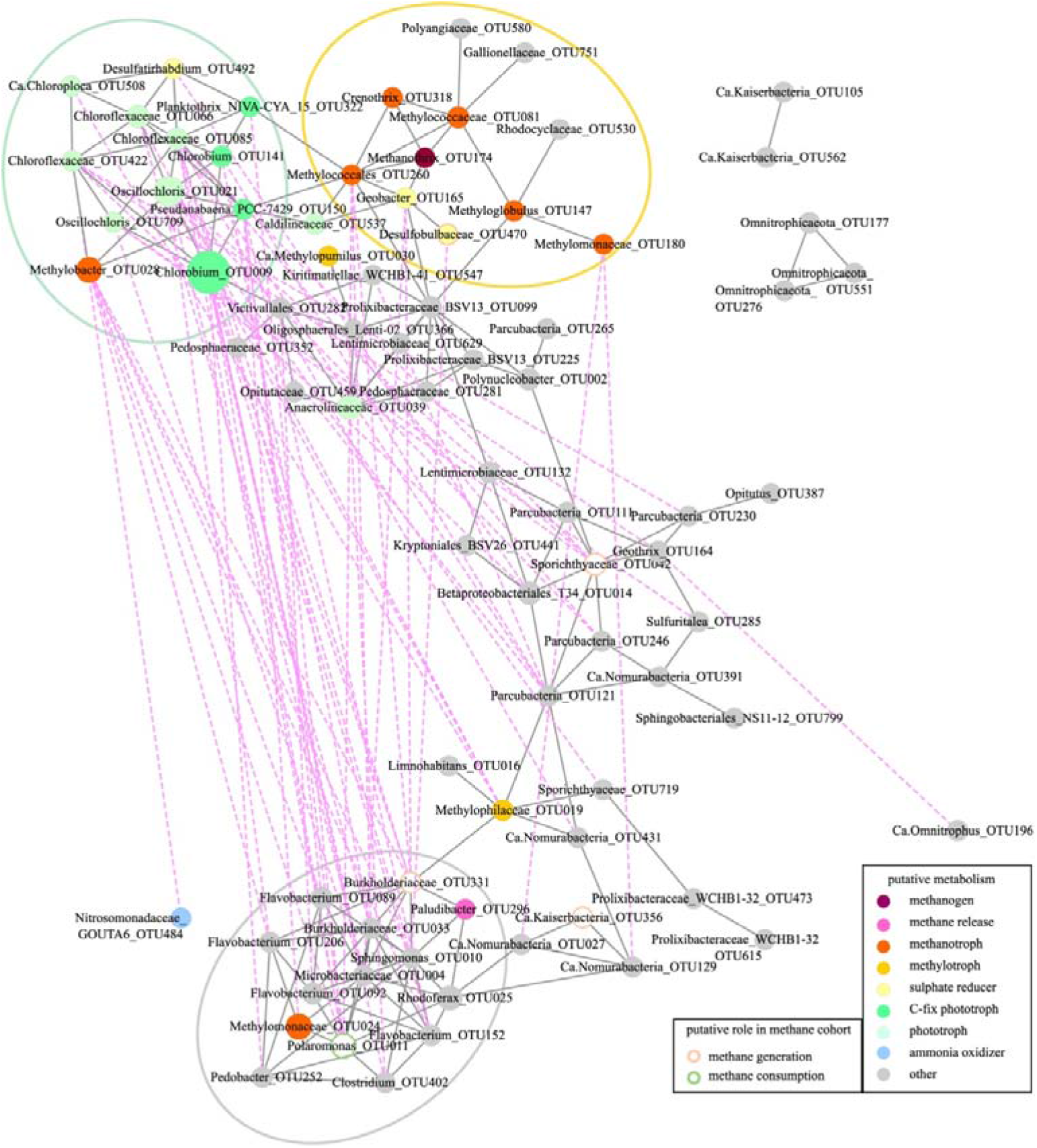
Lake time-depth-series network. Filled node-circles are OTUs with the following color codes: methanogens-red, methanotrophs-orange, methylotrophs-golden, C-fixing phototrophs-bright green, other phototrophs-pale green, sulphate reducers-yellow. Grey lines show positive correlations and pink-dotted lines show negative correlations. Green, golden, and grey bigger circles designate cohort clusters. We designated putative roles in methane consumption from observations in the dilution model communities and they are shown in not-filled node-circles in peach and green.

### Percentage of mixed cultures with growth varied with inoculum source

All inoculated cultures from the 2016 inoculum grew and produced sequences, while 71% of the cultures from 2017 sampling produced sequences. Despite similar culture conditions, there were distinct differences in the dominant genera in the cultures inoculated with water samples from early spring 2016 compared to autumn 2017 (Fig. 1D).

### Abundant lineages can both pass through and be retained by 0.22 μm filters

On average half of the top ten most abundant phylotypes in the time-depth-series and the 2017 culture-series were also detected in the media control sequenced from 2017. Specifically, five of the most prevalent phylotypes in the 2017 cultures (Pelomonas, Betaproteobacteriales - T34, Polynucleobacter, Sphingomonas, and Staphylococcus) were also detected in the 2017 media control. Seven of the top 25 most abundant OTUs in the time-depth series (Polaromonas, Polynucleobacter, Rhodoferax, Betaproteobacteriales - T34, an unclassified genus of the Sporichthyaceae family, Flavobacterium, and an unclassified genus of Burkholderiaceae) were abundant in media controls.

### Detection of methane cycling phylotypes does not always correlate to methane flux measurements

The majority (91%) of cultures had no detectable change in methane concentration or (60%) had no known methane cycling phylotypes (Figure 3A). Of note in our search for new organisms with a putative role in methane cycling were the 5% of cultures that had methane flux but no methane cycling phylotypes detected. Also of note were the 36 % of mixed cultures where obligate methane cycling phylotypes were detected but no methane flux was recorded. These two culture groups were noted and selected for further investigation.

**Figure 3.**
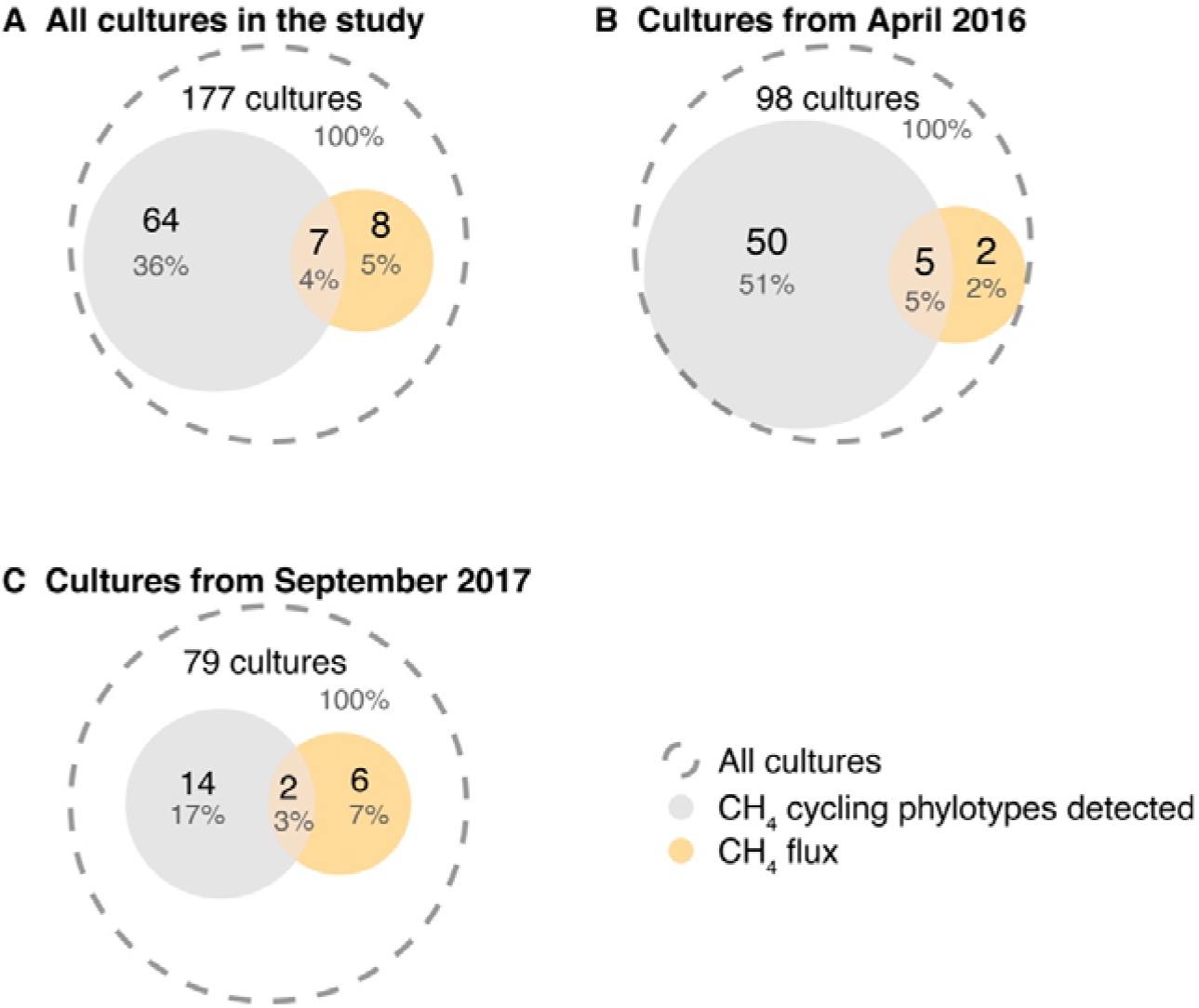
Venn diagram of all cultures in this study [A], cultures from April 2016 [B] and cultures from September 2017 [C]. In orange the cultures with measured methane flux and in grey the cultures with detection of (known) methane cycling phylotypes.

### Putative cohorts and potential newcomers to methane cycling

Due to the poor correlation between methane flux and methane cycling phylotypes, the co-occurrence patterns of the cultures of note were examined. Organisms detected uniquely in cultures that had unexplained methane concentration were recorded and a putative role in methane cycling proposed (Table 1). Eight microbes associated with the production of methane without detection of archaeal methanogens were noted by “CH4 production without methanogen”. One of these candidates, Rhodopseudomonas, is already annotated as a putative methane producer via the recently documented Fe-only nitrogenase methane release pathway (24, 25). Another cohort candidate, Desulfobulbaceae, is a putative sulphate reducer. Many cultures had abundant obligate methane consumers but no change in methane concentration, twenty-five microbes were uniquely associated with these cultures and were noted with “CH4 balance in the presence of methanotroph”. There were many Patescibacteria and one putative ammonia oxidizer, Nitrosomonadaceae, in this group. Six bacteria associated with consumption of methane without detection of methane consumers were noted by “CH4 consumption without CH4 consumer”. In total, 39 lineages were identified as potential contributors to methane cycling assemblages.

**Table 1.**
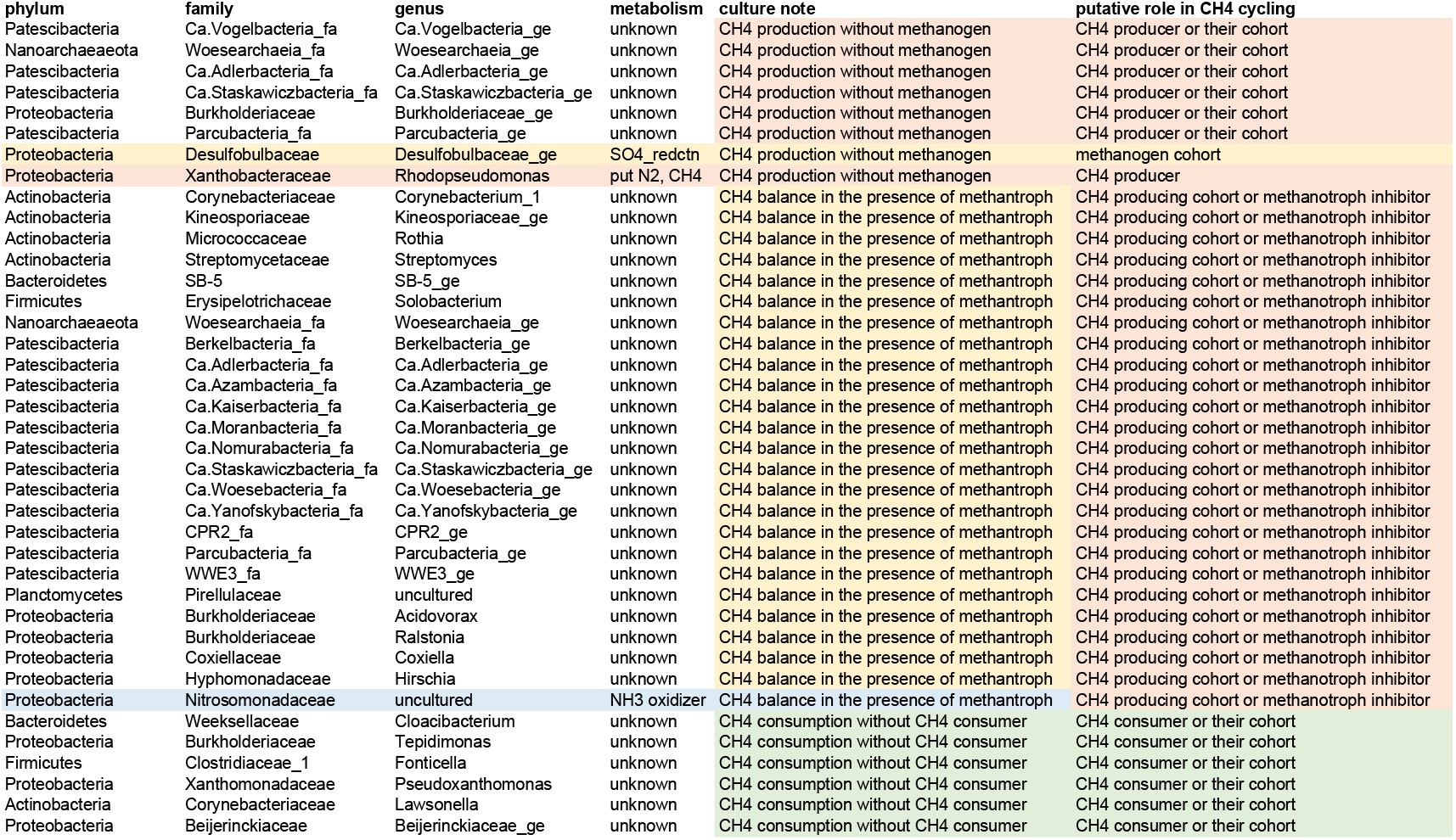
Phylotypes in this study with a potential role in methane cycling.

## Discussion

### Assembly of microbial communities is cell density and diversity dependent

The 71 % to 100 % success rate of culturing in this experiment is in stark contrast to the 10% to 30% success rate of dilution to extinction where only a few to maybe 20 cells are used in inoculant mixed cultures (11, 26). It therefore seems that around 50 cells of a natural community per inoculum (for anaerobic cultures from Lake Lomtjärnan) capture sufficient metabolic breadth to form a functional self-sustaining non-photic community, or cohorts. This for the first time places an upper limit on the minimum metabolic diversity required for functional (anaerobic) aquatic cohorts (27, 28).

### Filterable microbes play a significant role in natural and laboratory systems

A growing literature has discussed and evaluated the aquatic microbial filterable microbes in relation to the concept and efficacy of filter sterilisation of media used for laboratory culturing and environmental role (29–32). Here we also found that some phylotypes e.g., Polynucleobacter, can both pass through filters to be detected in filter ‘sterilised’ media, and be captured on filters as seen here in the time-depth-series. While the debate over whether filter sterilisation works is mostly complete, a discussion on the effect on abundance estimates of the loss of the filterable microbes might be worth re-opening.

### Associations between methanotrophs and other lifestyles in the lake

The lake time-series network shows an association between photoautotrophic Chlorobium, a methanotrophs, and sulphate reducing bacteria. This connection could be, for example, via carbon fixation by the autotrophic bacteria which also oxidise sulphur species to sulphate which is then consumed by the sulphate reducers. The Methylobacter phylotype, based on associations visualised in the network, is not reliant on proximity to a methanogen to thrive but rather is integral to the phototrophic cohort. The remainder of the methanotrophs in the cluster are associated with a methanogen and sulphate reducers. One of these sulphate reducers, Desulfobulbaceae is a putative methane consumer cohort member, while the other, Geobacter, is a known consort of Methanothrix (synonym Methanosaeta (40)) whereby direct electron transfer allows the methanogen to switch to CO2 utilisation for methane generation (44, 45). Methane is then released into the water and consumed by the cohort of methanotrophs.

Except for two cyanobacteria OTUs (Planktothrix and Pseudoanabaeana) most of the taxa in the green and gold circles are associated with oxic-anoxic interfaces. Besides the phototrophs all of the taxa present in the upper cohort and correlated to methanotrophs are potential microaerophiles e.g. Polyangjaceae or Gallionalaceae (33–35). Also included in the circles are potential anaerobe with known ability to live in microaerophile condition like Geobacter, or Rhodocyclaceae a taxa known to use a wide range of electron acceptor, including but not limited to O2 (36). These correlations between methanotrophs and taxa associated with the oxic anoxic interface is in line with the fact that this interface is a hotspot for methanotrophy (37–39). Based only on correlation it is impossible to tell if the cooccurrence of different OTU is due to shared environmental preference of by necessary interactions. The fact that all the organisms represented in the gold and green circles appear to favorize low oxygen environment suggests that the cooccurrence might be driven by environmental preferences. It is nevertheless interesting to observe that three different OTUs attributed to methanotrophic taxa are directly correlated to a methanogen, Methanothrix (synonym Methanosaeta (40)). The presence of an archaeal methanogen in a potentially oxic environment might seem surprising, but again there is also mounting evidence that methanogenesis isn’t limited to anoxic environments (41, 42). Furthermore Methanothrix is among the methanogen with a potential to strive in oxic conditions (42, 43). Interestingly though the Methanothrix OTU correlates with two OTUs that lean more on the anaerobic side of the interface (Geobacter and Crenothrix).

Others interesting correlation are the ones with iron cycling phylotypes (i.e Geobacter and Gallioneallaceae). Indeed both methanogen and methanotrophs have high iron demand for cofactor production (46). Furthermore, it has been suggested that interactions can be favorable to methanotrophs (47, 48). But if notable it is, again, impossible to clearly state the nature of the relation between those methane cycling taxa with the iron cycling ones.

All in all, the clusters in the top of the figure suggest that most of the methantrophs detected are attached to an environment with little oxygen favorizing taxa with the ability to deal with change in O2 concentration through mobility (like Gallionella or Chlorflexi) or metabolic plasticity (e.g Chloroflexi or Rhodocyladeae). Interestingly, the Methylococcales OTU abundances correlates with both anoxic leaning OTU (Chrenothrix, Methanothrix, Geobacter) and an oxygenic phototroph (Planktothrix and Pseudanabaena, both cyanobacteria). This again suggest microoxic condition, and is in line with work suggesting that aerobic methanotrophy can be enhanced by phototroph in low oxygen environment (49).

The most probable explanation for the negative correlation between the upper circle clusters (Figure 2) and the grey circle cluster in the bottom of the figure are due to spatial separation likely controlled by temperature. Several OTU present in the grey cluster belongs to taxa including both aerobic and anaerobic species like Polaromonas, Flavobacterium or Burkholdaerieacea. Both Polaromonas and Flavobacterium are generally considered psychrophilic and psychrotolerant taxa respectively (50, 51) (52). Furthermore Both Pedobacter and Rhodoferax also include psychrophilic taxa (53, 54). The association of these psychrophilic and tolerant taxa with the surface water, might be explained considering that sampling was performed during the ice-covered season it make sense as during winter, stratification is inverted, with the coldest water found at the surface directly beneath ice. It seems therefor very likely that the grey cluster is associated with cold temperature compared to rather than by oxygen concentration.

### Associations between methanotrophs and other lifestyles in the cohorts in model communities

Over the last two decades there has been a large increase in the catalogued number of microbes and metabolic pathways capable of generating, releasing, or consuming methane (55, 56). Both increases are likely to continue as we explore deeper genomic and metabolic space. For these reasons it is not a surprise that methane consumption and production did not correlate clearly with detection of known methane cycling phylotypes. It was surprising however, the degree to which these were decoupled. It therefore not clear if this uncoupling is due to poor flux detection, or low abundance of methane cycling organisms. Previously, it has been demonstrated that methanotroph abundance can be uncoupled form methane oxidation rates (58–60).

The generation here of a list of 39 novel phylotypes that may be directly or indirectly involved in methane cycling is a mere indicator of how much more work can be done to identify key players effecting cycling of methane and other C1 molecules.

We hypothesize that the participate in methane cycle is not only related to the presence of organisms with the known ability to produce or use methane, but that their cohorts as found in the model communities might be key controllers in methane cycling.

## Acknowledgments

SLG was supported by a SciLifeLab fellowship, a Stipend from King Carl XVI Gustav’s science foundation to research methane oxidation in lakes. SP was support by a SciLifeLab fellowship. Field, laboratory, and DNA amplicon sequencing was enabled by grants from Olsson-Borghs Stiftelse.

Computations were enabled by resources in projects 2020-5-529 & 2020-15-261 and data storage projects 2020-6-164 & 2020-16-196 provided by the Swedish National Infrastructure for Computing (SNIC) at UPPMAX which is partially funded by the Swedish Research Council through grant agreement no. 2018-05973.

SP, GM, SLG conceived of and implemented field and laboratory research. SP and SLG obtained financial and material resources while SP and RM obtained computational resources. RM and SLG wrote and implemented scripts for analysis and visualization of results. RM and SP drafted the manuscript and all authors contributed to discussions, editing, and approval of final manuscript. RM curated data and scripts. SP and SLG supervised and SLG coordinated the project.

